# Dog-wise canine gut metagenome assemblies with reconstructed bacterial genomes and viral candidates

**DOI:** 10.64898/2026.09.01.747583

**Authors:** Balázs Kakuk, Natheer Jameel Yaseen, Ákos Dörmő, Tamás Járay, Zsolt Boldogkői, Dóra Tombácz

## Abstract

Long-read metagenomic sequencing can improve genome recovery from complex gut microbial communities, yet directly reusable canine gut genome resources remain limited. Here we describe DogMAG, a canine gut metagenome resource based on dog-wise long-read and hybrid assemblies generated by grouping sequencing libraries according to canonical dog identity before assembly. The final dataset comprises 41 assemblies linked to 277 FASTQ records, including 30 Flye long-read-only and 11 OPERA-MS hybrid assemblies. A single integrated BASALT workflow produced 11,276 selected bin/version records, followed by explicit quality-based re-selection of 3,418 medium-quality-or-better metagenome-assembled genome candidates. External dRep dereplication yielded 792 strain-like representatives at 99% average nucleotide identity and 135 species/SGB-like representatives at 95%. GTDB-Tk classified all 792 representatives as Bacteria. Viral screening identified 22,068 geNomad predictions, of which 3,374 Complete, High-quality or Medium-quality viral/proviral candidate rows passed CheckV filtering with contamination ≤10%. DogMAG provides assemblies, genome and viral candidate sequences, metadata, provenance tables and workflow scripts for reuse, benchmarking and reanalysis.

## Background and Summary

The domestic dog (*Canis lupus familiaris*) is an important comparative model for microbiome research with relevance to veterinary medicine, translational host-microbiome studies and companion-animal health. Dogs share close environments, diets and microbial exposures with humans, while also developing naturally occurring diseases that can be studied in a clinically relevant non-laboratory setting [1–3]. Whole-genome metagenomic studies have shown that the canine gut microbiome shares functional similarities with the human gut microbiome, including diet-associated functional responses, but canine gut microbial genome resources remain less extensive than those available for humans [3].

Amplicon surveys, short-read shotgun metagenomes and genome-resolved canine or companion-animal resources have expanded knowledge of the dog gut microbiome [4–8]. DogMAG complements these resources by adding a directly reusable dog-wise canine gut metagenome assembly resource with linked BASALT-derived bacterial genome candidates, dereplicated MAG representative panels, viral/prophage candidates, metadata and workflow provenance. Its main contribution is the coordinated release of reusable data layers, including the BASALT-derived MAG candidate pool, strain-like and species/SGB-like MAG representative panels for mapping independent canine gut metagenomic reads and comparing read recruitment against alternative reference panels, GTDB-Tk taxonomy, viral/prophage candidate context, read-to-assembly provenance and workflow code. This structure allows users to map independent canine gut metagenomic reads to DogMAG genome panels, review candidate-selection criteria, rerun dereplication at alternative thresholds, update taxonomy and reannotate viral candidates as reference databases improve.

DogMAG uses canonical dog identity as the primary assembly unit. Sequencing project, date, diet, extraction batch, kennel, library preparation and run origin remain part of the metadata, but they are not the main axis of the assembly and binning workflow. This structure matches the biological unit used for genome recovery: reads from the same dog are assembled together where appropriate, and the resulting dog-wise assemblies are integrated in one BASALT run.

The dog-first assembly layer contains 41 final BASALT input assemblies: 30 Flye long-read-only dog-wise assemblies for dogs with long-read data and no retained short-read support for final hybrid assembly, and 11 OPERA-MS hybrid assemblies for dogs with both long-read and short-read data retained for the final BASALT run. These assemblies are linked to 277 FASTQ records, including 87 long-read FASTQ rows and 190 short-read FASTQ rows, in the public read-to-assembly manifest provided as Supplementary Table 4. The dog-first assembly design and final assembly-class composition are summarised in Figs. 1 and 2.

**Figure 1.**
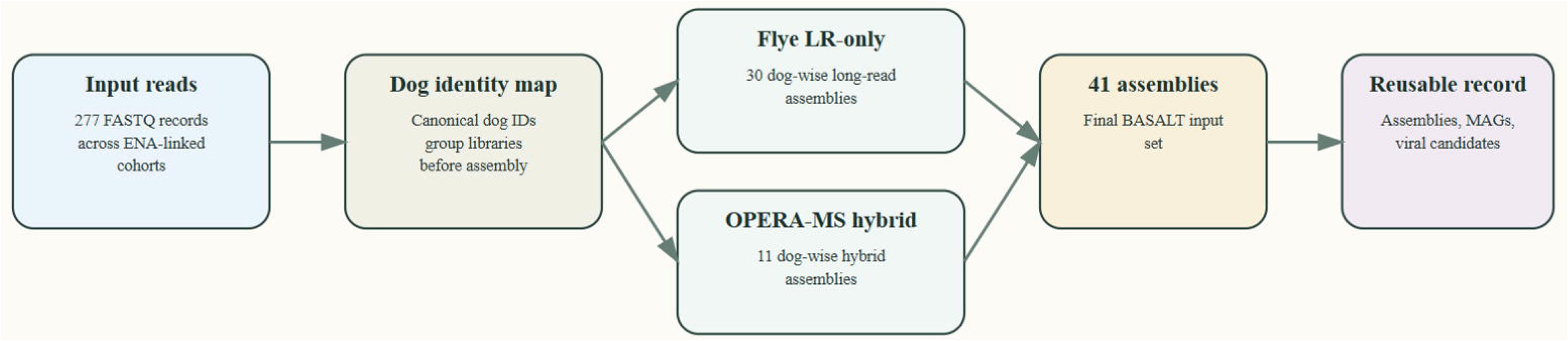
Dog-first assembly design and read-to-assembly provenance. Schematic of the DogMAG dog-first workflow. Sequencing libraries are grouped by canonical dog identity before assembly. Dogs with long-read data and no retained short-read support are assembled with Flye in metagenome mode, whereas dogs with retained long-read and short-read support are assembled with OPERA-MS. The final BASALT input set comprises 30 Flye long-read-only assemblies and 11 OPERA-MS hybrid assemblies linked to 277 FASTQ records in the read-to-assembly manifest.

**Figure 2.**
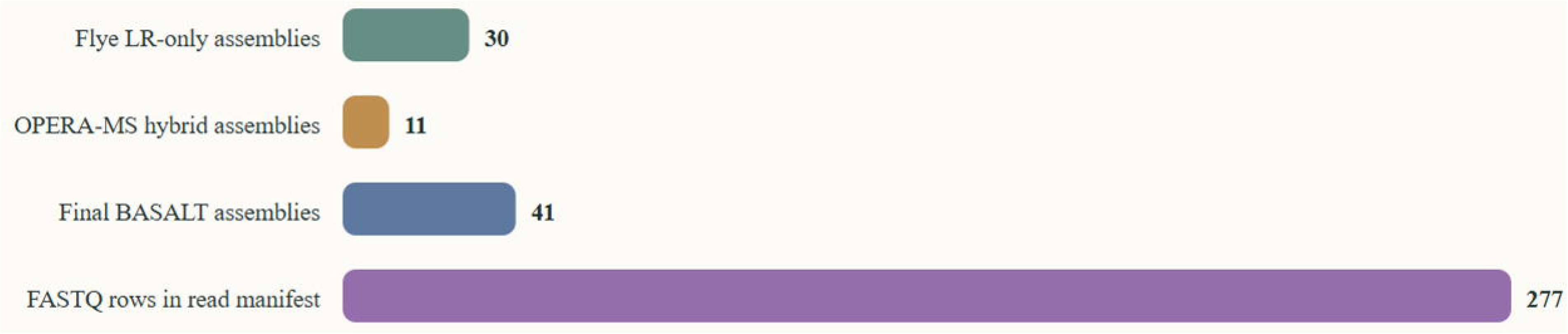
Final BASALT assembly classes and read-provenance coverage. Summary of the 41 dog-wise assemblies used as final BASALT inputs and the 277 FASTQ records linked to them in the read-to-assembly manifest. The final BASALT input layer contains 30 Flye long-read-only assemblies and 11 OPERA-MS hybrid assemblies. Counts are grouped by assembly class and manifest coverage rather than by historical project subset.

The dog-first BASALT workflow used manifest-driven assembly staging, read-to-assembly provenance tracking and CheckM2-backed quality estimation. BASALT was retained as the bin-comparison and BestBinset selection engine, while the dog-wise assembly design ensured that all retained final assemblies were processed in a single integrated run. The completed BASALT and candidate re-selection workflow produced 11,276 selected bin/version records from 30,556 polished or reassembled candidate versions. Applying explicit DogMAG quality thresholds and candidate re-selection retained 3,418 medium-quality-or-better MAG candidates for external ANI dereplication, including 503 high-completeness/low-contamination candidates and 2,915 medium-only candidates. External dRep dereplication of the reselected MAG candidate pool produced 792 99% ANI strain-like representatives and 135 95% ANI species/SGB-like representatives; the latter are not official SGB assignments. The BASALT-to-catalogue workflow and candidate attrition are summarised in Figs. 3 and 4.

**Figure 3.**
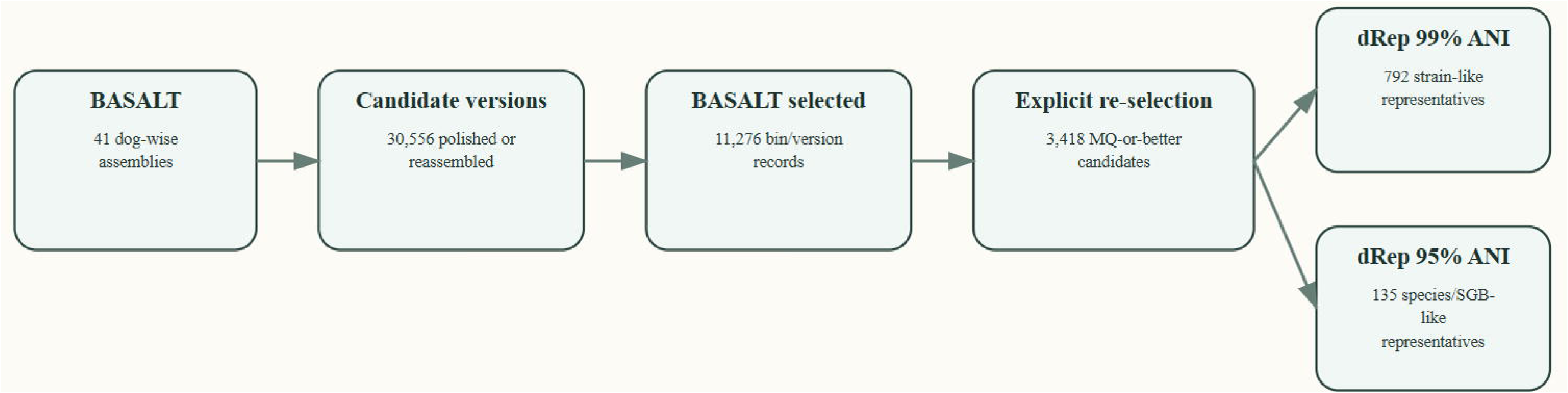
Dog-first BASALT workflow. Overview of BASALT workspace construction, assembly filtering, binner outputs, CheckM2 quality estimation, BASALT bin selection, long-read-supported bin context, candidate re-selection and final candidate-bin generation. The workflow selected 11,276 BASALT bin/version records from 30,556 candidate versions and retained 3,418 medium-quality-or-better MAG candidates after explicit completeness/contamination-based re-selection.

**Figure 4.**
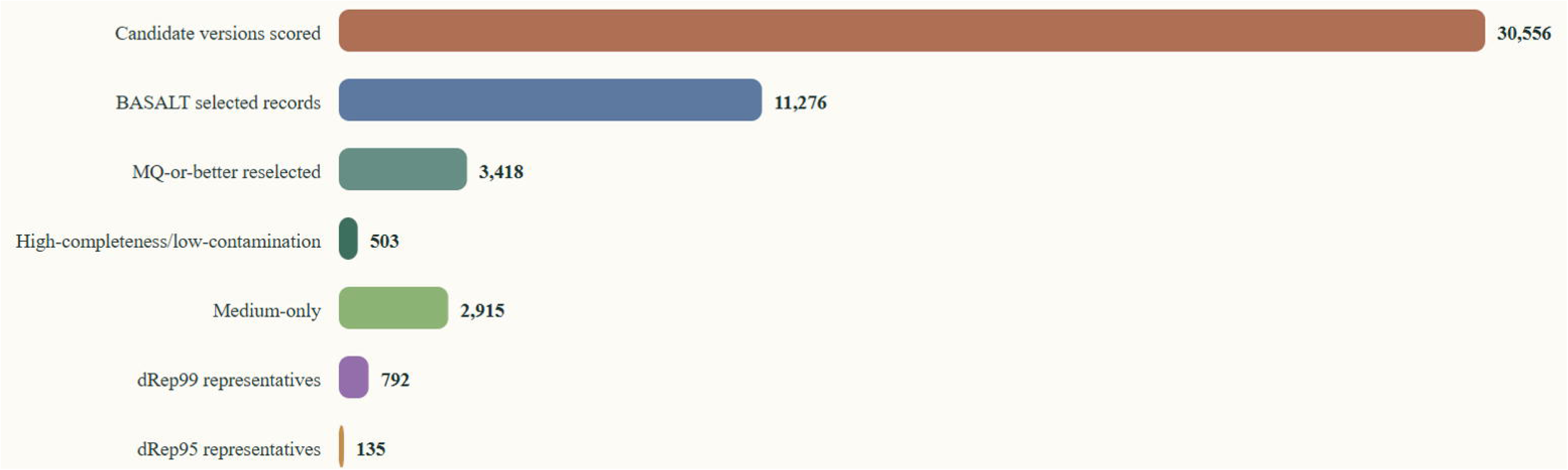
MAG candidate attrition and quality distribution. Attrition from 11,276 BASALT selected bin/version records and 30,556 polished/reassembled candidate versions to 3,418 medium-quality-or-better reselected MAG candidates, including 503 high-completeness/low-contamination candidates and 2,915 medium-only candidates. External catalogue-unit dRep dereplication yielded 792 99% ANI strain-like representatives and 135 95% ANI species/SGB-like representatives. Panel shows candidate attrition, operational quality classes and catalogue-unit dereplication counts for the BASALT-derived candidate pool and dereplicated representative sets.

GTDB-Tk classified all 792 strain-like representatives as Bacteria. Species labels were assigned for 768 representatives and genus labels for 791. The catalogue spans 132 unique species labels, 76 genera, 32 families, 20 orders, 11 classes and 7 phyla. Dominant phyla were Bacillota (449), Bacteroidota (120), Actinomycetota (74), Bacillota_I (50), Fusobacteriota (43), Pseudomonadota (42); rank-wise taxonomic composition is shown in Fig. 5. Viral screening identified 18,150 unbinned viral predictions and 3,918 binned/prophage-context viral predictions with geNomad, of which 2,306 unbinned and 1,068 binned/prophage candidates passed the final CheckV Complete/High/Medium and contamination <=10% filter.

**Figure 5.**
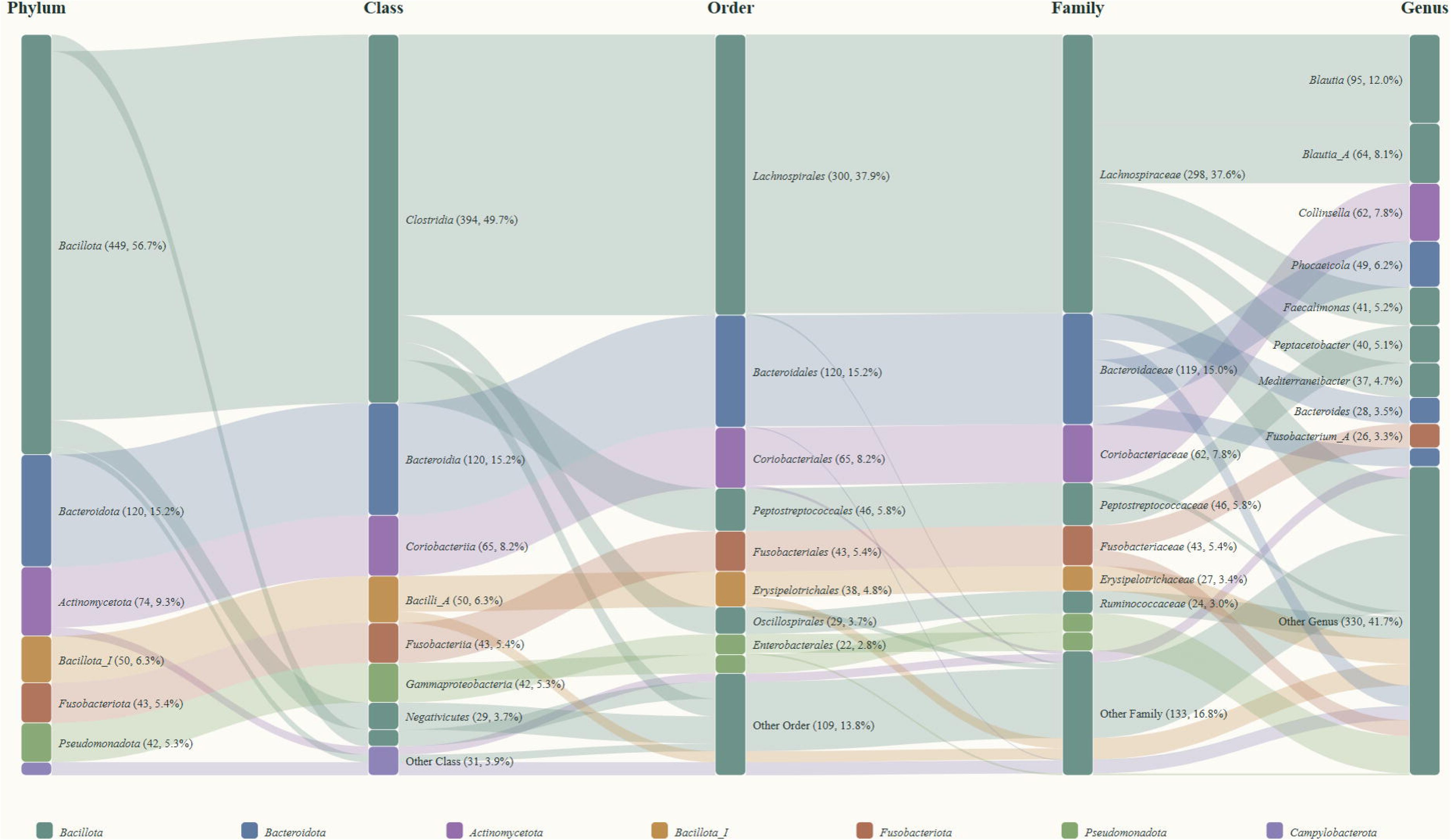
GTDB-Tk taxonomic composition of 792 dRep99 representatives. Descriptive alluvial summary of GTDB-Tk taxonomy for the 792 externally dereplicated 99% ANI representatives. All representatives were classified as Bacteria, with species labels assigned for 768 representatives and genus labels for 791. Dominant phyla were Bacillota (449), Bacteroidota (120), Actinomycetota (74), Bacillota_I (50), Fusobacteriota (43) and Pseudomonadota (42). Flows show genome counts from phylum to class, order, family and genus; long-tailed ranks are collapsed into rank-specific Other categories where needed, and colours follow the phylum-level assignment.

The primary reusable outputs of the DogMAG data record are: (1) a read-to-assembly provenance table linking public assembly identifiers to input read records; (2) filtered final dog-wise assembly FASTA files used as BASALT inputs; (3) assembly metrics for the dog-wise Flye and OPERA-MS assemblies; (4) summary-level BASALT workflow provenance and CheckM2-derived quality metadata; (5) metadata and FASTA files for the reselected medium-quality-or-better MAG candidate pool after operational completeness/contamination filtering; (6) final MAG representatives after explicit external ANI dereplication at 99% ANI and 95% ANI, with completeness, contamination, contiguity, taxonomy and provenance metadata; (7) GTDB-Tk genome taxonomy for the 99% ANI representative set; (8) viral and prophage candidate FASTA files and context tables regenerated from the dog-first assemblies and final BASALT bin context; and (9) workflow scripts and provenance notes documenting assembly generation, BASALT input construction, quality filtering, external dereplication and candidate-resource finalisation.

DogMAG is designed as a reusable Data Descriptor rather than as a hypothesis-testing study of canine gut microbiome biology. Age-, diet-, extraction- and method-associated biological analyses using overlapping sample sets are reported in previous or companion studies and are not treated as conclusions of this manuscript. The present manuscript describes the assembly products, genome-resolved candidate resource, viral/prophage candidate layer and the workflow provenance needed for independent reuse.

As an additional technical validation of reference-panel coverage, 23 independent Waltham canine gut ONT metagenomes [8] and 23 mixed-kennel source-cohort libraries were mapped against bacterial RefSeq and the 135-member DogMAG 95% ANI panel with matched read-accounting settings. Across the 46 paired libraries, the median primary-read recruitment fraction was 61.07% with RefSeq and 79.07% with DogMAG; the corresponding median final assigned-read fractions were 60.21% and 78.88%. In the independent Waltham cohort, median recruitment increased from 51.02% to 75.73% and median final assignment increased from 49.84% to 75.48%.

## Methods

### Study design and sample provenance

DogMAG integrates canine faecal metagenomic sequencing data from related source studies and sequencing campaigns. In the current dog-first design, the primary workflow unit is the canonical dog identity rather than the historical project subset. Source project labels remain in metadata fields because they document sampling provenance, sequencing platform, wet-lab protocol, publication status and ENA accession origin, but they are not used to split the final BASALT workflow into separate project-specific runs.

Sample-linked input reads used for DogMAG assembly and MAG generation are identified by ENA project accessions PRJEB75753 for Dog_M0 input data, PRJEB82125 for DMD input and provenance records, PRJEB85420 for CaniMeta/Serteperti input and provenance records, and PRJEB115259 for DogMAG sample registrations, newly deposited mixed-kennel long-read WGS data and associated DogMAG genome records. Dog_M0 and Serteperti source-data provenance is described in the associated Scientific Data descriptor [24]. Previously generated reads are cited here to document sample provenance and input-read availability; the DogMAG data record reports dog-wise assemblies, BASALT workflow summaries and reselected MAG candidate metadata, viral/prophage candidates and workflow metadata generated from these inputs.

The read-to-assembly provenance data are provided in Supplementary Table 4, which records one row per linked FASTQ using stable public assembly, sample and read identifiers.

### DNA isolation, library preparation and sequencing

The source sequencing datasets were generated with multiple wet-lab workflows. DNA isolation, library preparation and sequencing provenance are therefore recorded at the library level rather than described as a single universal protocol. Supplementary Table 3 records source dataset, public sample identifier, sequencing platform, library name, extraction workflow, library preparation workflow, DNA concentration, concentration unit and source metadata provenance for each included sequencing library.

For newly submitted mixed-kennel long-read WGS libraries, the ENA metadata records PromethION sequencing and ZymoBIOMICS MagBead DNA/RNA extraction from canine faecal samples. These libraries were prepared as Oxford Nanopore native-barcoded ligation libraries and sequenced on PromethION R10 flow cells. Basecalling was performed with Dorado v0.8.3 using the super-accurate model, followed by barcode demultiplexing. This basecalling description applies specifically to the newly submitted mixed-kennel PromethION libraries; DMD, Serteperti and previously published source datasets are documented through their source records and publications.

For DMD long-read WGS libraries, the ENA template records ONT MinION sequencing. Extraction method is encoded in the library names, including MN for Macherey-Nagel preparations and Zymo_HMW for Zymo Research Quick-DNA HMW MagBead preparations. These libraries were prepared as native-barcoded ONT ligation libraries and sequenced on MinION.

For Serteperti long-read WGS libraries, the ENA template records ONT MinION sequencing. Extraction method is encoded as HMW or ZymoBIOMICS_96_MagBead, corresponding to Quick-DNA HMW MagBead and ZymoBIOMICS 96 MagBead DNA extraction workflows, respectively. These libraries were prepared as ONT long-read WGS libraries and sequenced on MinION.

Previously published Dog_M0/Toti and method-comparison sequencing datasets are documented through their source publications. In the Scientific Data canine faecal microbiome dataset [24], Dog_M0/Toti was a single adult dog faecal sample collected shortly after defecation, stored initially at -20 degrees C and transferred to -80 degrees C within 24 h. DNA extraction, Illumina NovaSeq paired-end sequencing and ONT MinION long-read sequencing details are cited from that source rather than restated as new wet-lab work.

### Dog-first assembly strategy

The dog-first assembly strategy grouped sequencing libraries by canonical dog identity before assembly. Sequencing project, date, diet, extraction batch, kennel, library preparation, sequencing run and project origin were retained as metadata fields, but they were not used to split the final dog-level assembly units. This design was chosen because the individual dog was the biological unit for downstream genome recovery.

Dogs with long-read data but no retained paired short-read support were assembled with Flye in metagenome mode [9]. Depending on the retained assembly batch, Flye v2.9.4-b1799 or v2.9.6-b1802 was run with --meta and --nano-hq. Dogs with both retained short-read and long-read libraries were assembled with OPERA-MS [10]. The OPERA-MS workflow used the local OPERA-MS installation corresponding to Git revision 026f9a5f58ba6dca626f677bb50521fa423901ec, with minimap2 v2.24-r1122, SAMtools/HTSlib v1.23.1 and NOPOLISHING=YES.

The final BASALT input set contained 41 dog-wise assemblies: 30 Flye long-read-only assemblies and 11 OPERA-MS hybrid assemblies. Alternative hybrid assemblies generated with metaSPAdes or XelNaga were produced during method exploration but were not included in the final BASALT input set. Public assembly-to-read relationships are provided in Supplementary Table 4, and the workflow scripts used for Flye assembly, OPERA-MS assembly and assembly collection are available in the DogMAG repository.

### Assembly preparation for BASALT

Final dog-wise assemblies were collected into a merged BASALT input layer and filtered with a minimum contig-length threshold of 1,500 bp. The retained input set comprised the same 41 assemblies described above. Assembly-level sequence metrics for these retained inputs, including total assembly length, contig count, N50, N90, longest contig, GC percentage, linked FASTQ-record count, ENA coassembly BioSample accession and ERZ analysis accession, are provided in Supplementary Table 8.

### BASALT input construction

The final dog-first BASALT workspace contained the assembly links, depth matrices, binner outputs and metadata used for the integrated BASALT run. BASALT input files were generated from the dog-first assembly and read manifests, retaining Flye long-read-only and OPERA-MS hybrid assemblies while excluding exploratory alternative assembler outputs. Full command lines, script names and workflow-provenance details are provided in the DogMAG repository and associated article data package.

### BASALT execution

Genome binning and bin selection were performed with the DogMAG BASALT v1.2.0 fork [11]. The DogMAG production workflow was based on upstream BASALT commit 5f51ba3780749be2fbb7c868df505548f894f7e4, with local modifications to improve restart safety, CheckM2-output validation, storage handling and execution on large datasets. A cleaned, publication-oriented representation of these generalized modifications, excluding dataset-specific rescue scripts and checkpoint edits, is archived in the Balays/BASALT fork as the DogMAG source snapshot at commit bdd4106a398db69bee83ea9ad1d331b00c041122.

The integrated workflow used MetaBAT2 v2.18, MaxBin2 v2.2.7, CONCOCT v1.1.0, SemiBin2 v2.2.1 and single-contig binning outputs where available [12–15]. MetaBAT2 bins were generated across minimum-contig settings of 200, 300, 400 and 500 bp with --maxEdges 200; MaxBin2 used probability thresholds of 0.3, 0.5, 0.7 and 0.9; CONCOCT used cluster-count settings of 100 and 200; and SemiBin2 was run with single_easy_bin and --sequencing-type=long_read. BASALT was used as the central bin-comparison and BestBinset selection engine. Quality estimation used CheckM2 v1.1.0 with the uniref100.KO.1.dmnd database [16].

The final BASALT run processed the 41 dog-wise assemblies together with their linked short-read and long-read inputs using -q checkm2, --mode continue, --module all, output prefix dogfirst, 28 threads and 300 GB memory. Full command lines and workflow-provenance details are provided in the DogMAG repository and associated article data package.

BASALT generated per-assembly binner outputs, selected best-bin sets, quality reports, long-read-supported bin context and candidate bin/MAG metadata. Completeness and contamination estimates used in the DogMAG bin metadata originate from the CheckM2 quality reports generated during the BASALT workflow. The completed BASALT bin-selection stage produced 11,276 selected bin/version records, which were then filtered with the operational DogMAG completeness and contamination thresholds described below.

### MAG candidate quality filtering and external dereplication

BASALT produced 11,276 selected bin/version records from 30,556 polished or reassembled candidate versions. Because BASALT’s internal comparison and redundancy handling is not equivalent to a fixed external catalogue-unit definition, DogMAG applies an explicit post-BASALT candidate re-selection and external dereplication workflow before reporting catalogue-level counts.

For each original BASALT bin group, all available polished and reassembled candidate versions were compared using CheckM2-derived completeness and contamination estimates [16]. Candidate versions were considered eligible for the medium-quality-or-better catalogue pool when completeness was >= 50% and contamination was <= 10%, following operational MIMAG-style thresholds [20]. High-completeness/low-contamination candidates were defined as completeness >= 90% and contamination <= 5%. When multiple eligible versions were available for the same original bin group, candidates were ranked using the quality score completeness - 5 × contamination. N50 and related contiguity fields were used as tie-breakers where implemented in the re-selection script. This candidate re-selection retained 3,418 medium-quality-or-better MAG candidates, including 503 high-completeness/low-contamination candidates and 2,915 medium-only candidates. A further 7,858 original bin groups did not have an eligible medium-quality candidate.

The reselected MAG candidate pool was externally dereplicated with dRep v3.6.2 [17] using 52 processors, minimum genome length 50,000 bp, completeness >= 50% and contamination <= 10%. Mash primary clustering used a sketch size of 1,000, followed by fastANI secondary comparisons [18,19]. The 95% ANI run used primary and secondary ANI thresholds of 0.90 and 0.95, respectively; the 99% ANI run used thresholds of 0.95 and 0.99. Both runs used a minimum comparison coverage of 0.10, the larger-genome coverage denominator, average-linkage clustering and default representative scoring weights: completeness 1, contamination 5, strain heterogeneity 1, N50 0.5 and centrality 1. The workflow completed 172,570 fastANI comparisons at 95% ANI and 164,422 at 99% ANI. dRep’s internal CheckM-based filtering retained 3,222 genomes, corresponding to 94.27% of the reselected pool, before final representative selection. The final outputs contained 792 99% ANI strain-like representatives and 135 95% ANI species/SGB-like representatives.

The 99% and 95% ANI representative counts are external dereplication layers applied after BASALT, not BASALT-internal BestBinset counts. The 95% ANI units are described as species/SGB-like representatives because the clustering threshold approximates a species-level genome catalogue unit, but the workflow does not reproduce official SGB construction or placement procedures used by other catalogues.

### Prokaryotic genome taxonomy

Taxonomic classification was performed on the 792 externally dereplicated 99% ANI representatives using GTDB-Tk v2.7.1 with GTDB release r232 [21]. GTDB-Tk was run after candidate re-selection and dRep dereplication so that taxonomy describes the final strain-like representative set rather than all pre-dereplication BASALT candidates.

All 792 representatives were classified as Bacteria and no genomes were excluded from the GTDB-Tk summary. GTDB-Tk assigned species labels to 768 representatives and genus labels to 791. Across the representative set, GTDB-Tk reported 132 unique species labels, 76 genera, 32 families, 20 orders, 11 classes and 7 phyla. Classification used ANI screening for 768 representatives, topology-defined placement for 23 representatives and RED-based novelty assessment for 1 representative. The median closest GTDB ANI was 98.04% and the median alignment fraction was 0.827.

### Reference-panel read-recruitment comparison

The technical coverage of the DogMAG species/SGB-like reference layer was evaluated by mapping an equal-sized set of ONT long-read metagenomes against DogMAG and bacterial RefSeq. The balanced comparison comprised 23 libraries from the mixed-kennel source cohort and 23 independent Waltham canine gut metagenomes. The Waltham libraries were not used to construct DogMAG and therefore provide an external read-recruitment assessment, whereas the mixed-kennel libraries provide a source-cohort benchmark. The same FASTQ files were used for both reference panels. For the RefSeq arm, previously completed mixed-kennel alignments were reused after confirming the original FASTQ identities, reference index and mapping parameters; the 23 Waltham libraries were mapped de novo. All 46 libraries were mapped de novo against the 135-member DogMAG 95% ANI panel.

Read mapping and taxonomic assignment were performed with minitax v0.9 [4] using the ONT map-ont preset and the mm2-fast minimap2-compatible mapper. Both arms retained up to 20 secondary alignments per read and applied the same post-alignment settings: selection of the highest-MAPQ candidates, retention of alignments with the highest CIGAR score and taxonomic refinement with the BestAln method at a threshold of 0.6. SpeciesEstimate summaries were generated as an additional deposited output. The RefSeq panel used a bacterial RefSeq minimap2 index and matched NCBI taxonomy tables; the DogMAG panel used the final 95% ANI DogMAG reference index and the GTDB-Tk-derived taxonomy table. Primary-read accounting was calculated separately from secondary-alignment counts. For each library and panel, the reported validation fields comprise primary reads examined, primary reads mapped, mapped-read fraction, reads receiving a final BestAln taxonomic assignment and assigned-read fraction. Comparisons were paired by library and summarised separately for the mixed-kennel source cohort and independent Waltham cohort using per-library medians and interquartile ranges; pooled read-weighted fractions were also calculated from summed primary-read counts. The consistency of the paired direction of change was assessed with a two-sided exact binomial sign test after excluding exact ties.

### Viral and prophage genome candidates

Viral and prophage candidates were recovered with a two-branch workflow that separates unbinned/free-virus discovery from binned/prophage-context discovery. geNomad v1.12.0 with database v1.9 [22] was used for viral and plasmid prediction, and CheckV v1.1.1 with database v1.5 [23] was used for viral candidate quality estimation. The unbinned branch analysed contigs from the final dog-first assembly layer. The binned/prophage-context branch analysed contigs associated with BASALT-derived bin/MAG context so that candidate viral sequences could be linked back to genome bins where appropriate.

geNomad predicted 18,150 unbinned viral sequences and 3,918 binned/prophage-context viral sequences, for 22,068 viral predictions in total. geNomad also reported 14,739 unbinned plasmid predictions and 6,052 binned plasmid predictions; these plasmid predictions are tracked separately and are not counted as viral/proviral candidates in the final viral summary. After CheckV filtering to Complete, High-quality or Medium-quality candidates with contamination <= 10%, 2,306 unbinned and 1,068 binned/prophage-context viral candidate rows remained, for 3,374 final filtered viral/proviral candidate rows. These rows corresponded to 3,282 unique source-contig MD5s, indicating that a small number of source contigs were observed in more than one context.

The final viral/prophage results are reported as candidate viral or proviral sequences, not as a dereplicated viral operational taxonomic unit (vOTU) catalogue. Viral dereplication is intentionally deferred so that users can apply their preferred vOTU thresholds and tools to the deposited sequence and metadata tables. Broad viral taxonomy was dominated by Caudoviricetes in both unbinned and binned/prophage-context branches, based on geNomad taxonomic assignments.

### Data Records

The DogMAG-derived dataset is organised for direct reuse through ENA and an associated article data package. The deposited resource comprises dog-first assemblies, read-to-assembly provenance, summary-level BASALT provenance and quality metadata, the reselected medium-quality-or-better MAG candidate pool, externally dereplicated MAG representative layers, viral/prophage candidate records and workflow scripts. The complete BASALT workspace, transient intermediate files and the full set of polished or reassembled candidate versions are not part of the deposited resource.

The associated article data package is organised into metadata, assemblies, BASALT-derived outputs, MAG candidates and representatives, viral/prophage candidates, reference-panel validation outputs, workflow documentation and scripts. Key metadata tables include sample, library and read metadata, the read-to-assembly manifest, assembly metrics and checksums, BASALT candidate and quality summaries, MAG metadata and quality summaries, ANI95 and ANI99 cluster tables, GTDB-Tk taxonomy, viral candidate context tables and reference-panel read-accounting summaries. Full command lines, software versions and workflow notes are provided in the workflow documentation.

Supplementary Table 4 links each public dog-wise assembly identifier to public read and sample identifiers, read role, sequencing platform, ENA study accession and checksum fields where available. The manuscript Methods, public workflow scripts and supplementary provenance tables record dog-wise assembly generation, BASALT input construction, CheckM2 use, operational quality filtering and downstream catalogue finalisation.

Raw sequencing reads, primary metagenome assemblies, derived MAG BioSamples and MAG assembly records have been deposited in the European Nucleotide Archive (ENA) at EMBL-EBI under study accession PRJEB115259. The raw sequencing sample records comprise 84 faecal metagenome samples registered under ERS30645926-ERS30646009. The 41 dog-wise primary metagenome assemblies are linked to coassembly BioSamples ERS31153171-ERS31153211 and were accepted as ENA assembly analysis records ERZ29880033-ERZ29880073; assembly-level accessions, BASALT-derived coverage estimates and coverage provenance are listed in Supplementary Table 8. The derived MAG BioSample records comprise 135 DogMAG ANI95 representatives registered under ERS31049448-ERS31049582. Of the 135 DogMAG ANI95 representative MAGs, 122 multi-contig MAG assemblies were accepted by ENA as genome assembly analysis records and assigned ERZ accessions. These ERZ accessions, together with the retained data-package status of the remaining 13 single-contig representatives, are listed in Supplementary Table 9.

### Data Overview

The DogMAG data resource consists of five linked layers: input read provenance, dog-wise assemblies, BASALT-derived MAG candidate records, externally dereplicated MAG representatives and viral/prophage candidate records.

**Read provenance. The read-to-assembly manifest links 277 FASTQ records to the 41 final dog-wise BASALT input assemblies. The public supplementary table uses stable public sample and assembly identifiers.**

**Dog-wise assemblies.** The final BASALT input assembly set contains 30 Flye long-read-only assemblies and 11 OPERA-MS hybrid assemblies filtered at a minimum contig length of 1,500 bp. All 41 assemblies are deposited under PRJEB115259 as primary metagenome assembly analysis records ERZ29880033-ERZ29880073 and are linked to coassembly BioSamples ERS31153171-ERS31153211 (Supplementary Table 8).

**BASALT workflow summary and reselected candidates.** BASALT selected 11,276 bin/version records, and the internal final comparison table contained 30,556 polished or reassembled candidate versions. These values document workflow attrition. The reusable pre-dereplication genome layer comprises the 3,418 reselected medium-quality-or-better candidates reported in Supplementary Table 2; the complete BASALT workspace and all transient candidate-version files are not deposited.

**Reselected and externally dereplicated MAGs.** Explicit completeness/contamination-based candidate re-selection retained 3,418 medium-quality-or-better MAG candidates. External dRep dereplication yielded 792 99% ANI strain-like representatives and 135 95% ANI species/SGB-like representatives. GTDB-Tk taxonomy is provided for all 792 strain-like representatives.

**Viral/prophage candidates.** geNomad and CheckV outputs are provided for unbinned/free-virus candidates and binned/prophage-context candidates. The final contamination-filtered Complete/High/Medium set contains 3,374 viral/proviral candidate rows representing 3,282 unique source-contig MD5s.

The main article reports the 99% ANI representative set as the DogMAG strain-like MAG catalogue comparison layer and the 95% ANI set as a species/SGB-like comparison layer. The full pre-dereplication candidate pool remains available for users who want to rerun dereplication, compare scoring schemes or recover candidates under alternative thresholds.

## Technical Validation

### Assembly and MAG catalogue consistency

The final assembly layer comprises 41 dog-wise BASALT input assemblies linked to 277 FASTQ records in Supplementary Table 4: 30 Flye long-read-only assemblies and 11 OPERA-MS hybrid long-read/short-read assemblies. The same 41 public assembly identifiers are represented in the BASALT assembly index, final input list and ENA primary metagenome analysis records. All 41 Webin-CLI submissions passed, yielding ERZ29880033-ERZ29880073. Supplementary Table 8 links each assembly to its coassembly BioSample, ERZ accession, sequence metrics and length-weighted BASALT total-average-depth estimate.

The completed BASALT result contained 11,276 selected bin/version records and a comparison table with 30,556 polished or reassembled candidate versions. Candidate re-selection evaluated all 30,556 candidate rows across 11,276 original bin groups and retained 3,418 medium-quality-or-better groups. The remaining 7,858 groups did not contain an eligible candidate under the DogMAG completeness and contamination thresholds. Among retained groups, 503 candidates met the high-completeness/low-contamination threshold and 2,915 were medium-only.

External dRep dereplication was run on the reselected candidate FASTA set. dRep retained 3,222 genomes after its internal quality filtering, corresponding to 94.27% of the reselected pool. The 99% ANI run produced 792 strain-like representatives and the 95% ANI run produced 135 species/SGB-like representatives. These external dereplication outputs are reported as catalogue-comparison layers, separately from BASALT’s internal redundancy handling and from the raw 11,276 selected bin/version records.

### Genome taxonomy coverage

GTDB-Tk validation confirmed that all 792 99% ANI representatives were classified as Bacteria. No genomes were present in the GTDB-Tk excluded-genomes summary. Rank-level completeness was high: all 792 representatives received phylum-, class-, order- and family-level assignments, 791 received genus assignments and 768 received species labels. The dominant phyla were Bacillota (449), Bacteroidota (120), Actinomycetota (74), Bacillota_I (50), Fusobacteriota (43), Pseudomonadota (42). The most common genera included Blautia (95), Blautia_A (64), Collinsella (62), Phocaeicola (49), Faecalimonas (41), Peptacetobacter (40). The most common species labels included Blautia hansenii (42), Collinsella intestinalis (40), Blautia sp900556555 (36), Blautia_A sp900541345 (34) and Mediterraneibacter gnavus (30). These summaries are descriptive taxonomy results for the final 99% ANI representative set and are release-specific to the GTDB-Tk and representative-selection versions used here.

### Independent reference-panel read recruitment

Read recruitment against the DogMAG 95% ANI representative panel was evaluated relative to bacterial RefSeq using identical ONT mapping and taxonomic-assignment parameters implemented in minitax [4]. The balanced validation set comprised 23 long-read libraries from the mixed-kennel source cohort and 23 independent Waltham canine gut metagenomes; the latter provide an external assessment because they did not contribute to construction of the DogMAG catalogue. The two mapping arms contained identical primary-read denominators for all 46 libraries, representing 28,831,616 primary reads in total, and no library was missing from either arm.

Across all libraries, the median fraction of primary reads recruited was 61.07% for bacterial RefSeq (interquartile range (IQR), 47.96-70.98%) and 79.07% for DogMAG (IQR, 73.79-84.23%), corresponding to a median paired increase of 18.42 percentage points. DogMAG recruited a higher fraction in 44 of 46 libraries (two-sided exact sign-test P = 3.08e-11). The pooled read-weighted recruitment fractions were 55.03% and 75.28%, respectively, equivalent to 5,840,706 additional primary reads recruited by DogMAG under the matched workflow. Final BestAln taxonomic assignments showed a similar pattern: median assigned fractions were 60.21% for RefSeq (IQR, 46.54-70.09%) and 78.88% for DogMAG (IQR, 73.35-84.00%), corresponding to a median paired increase of 19.08 percentage points. DogMAG was higher in 45 of 46 libraries (P = 1.34e-12), and the pooled analysis contained 6,062,332 additional assigned reads.

The independent Waltham cohort showed the clearest gain. Median primary-read recruitment increased from 51.02% with RefSeq (IQR, 46.41-62.20%) to 75.73% with DogMAG (IQR, 72.66-79.58%), with a median paired increase of 23.89 percentage points; DogMAG was higher for all 23 Waltham libraries (P = 2.38e-7). Median final assignment increased from 49.84% to 75.48%, with a median paired increase of 24.59 percentage points. In the mixed-kennel source cohort, median recruitment increased from 69.90% to 84.32% (median paired increase, 15.44 percentage points), and median final assignment increased from 69.08% to 84.09% (median paired increase, 15.85 percentage points). Among mapped primary reads, the median fraction receiving a final assignment was 98.32% for RefSeq and 99.65% for DogMAG. These results support higher canine gut read recruitment and final assignment by the DogMAG panel in this matched comparison (Fig. 6); they are not interpreted as biological differences between cohorts or as direct comparisons of database-specific community composition.

**Figure 6.**
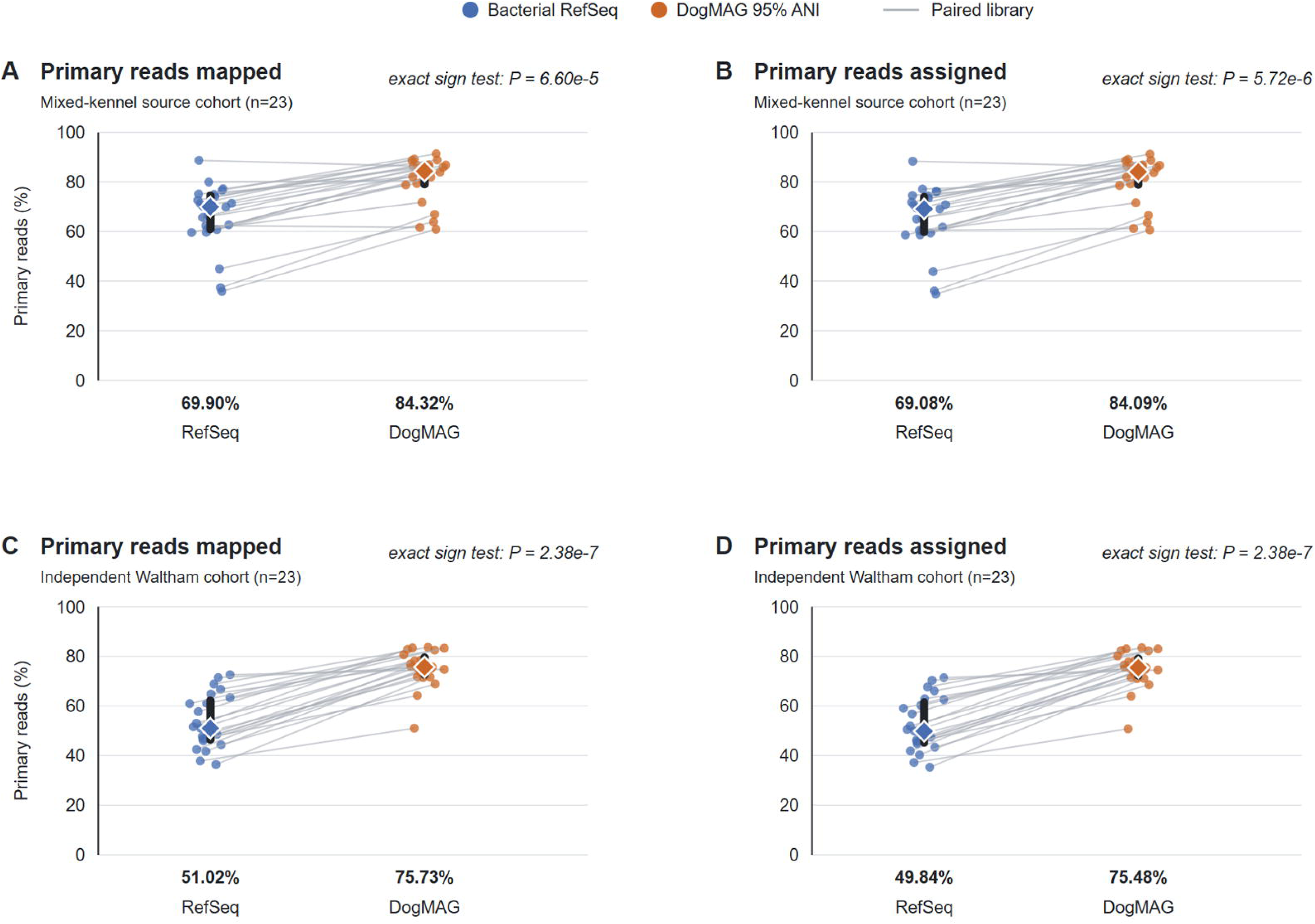
DogMAG read recruitment and taxonomic assignment relative to bacterial RefSeq. Paired primary-read recruitment and final assigned-read fractions for 23 mixed-kennel source-cohort libraries and 23 independent Waltham canine gut ONT metagenomes analysed against bacterial RefSeq and the 135-member DogMAG 95% ANI representative panel using minitax. Points represent individual libraries and grey lines connect results obtained from the same library against the two reference panels. Large diamonds show cohort medians and thick vertical bars show interquartile ranges. Median values are printed below each reference panel. P values are from two-sided exact binomial sign tests of the paired direction of change after excluding exact ties. In the mixed-kennel source cohort, median recruitment increased from 69.90% to 84.32% and median final assignment increased from 69.08% to 84.09%. In the independent Waltham cohort, median recruitment increased from 51.02% to 75.73% and median final assignment increased from 49.84% to 75.48%.

### Viral/prophage candidate quality control

Viral validation used geNomad prediction followed by CheckV quality estimation and contamination filtering. The unbinned branch produced 18,150 geNomad viral predictions and the binned/prophage-context branch produced 3,918, for 22,068 viral predictions in total. After retaining only CheckV Complete, High-quality or Medium-quality candidates with contamination <= 10%, 2,306 unbinned and 1,068 binned/prophage-context candidate rows remained. The combined final filtered table contained 3,374 rows, 3,282 unique source-contig MD5s, median length 39,364 bp, median CheckV completeness 87.14% and median CheckV contamination 0.00%.

The final filtered viral candidates were Caudoviricetes-dominated at the class-like geNomad taxonomy field (Caudoviricetes (3,299), Malgrandaviricetes (51), Arfiviricetes (8), Quintoviricetes (6), unclassified (5), Faserviricetes (3) and Revtraviricetes (2)). These class-like assignments account for all 3,374 final filtered candidate rows. Because viral dereplication was not performed, these counts describe candidate viral/proviral sequences and source-contig contexts rather than nonredundant vOTUs (Fig. 7).

**Figure 7.**
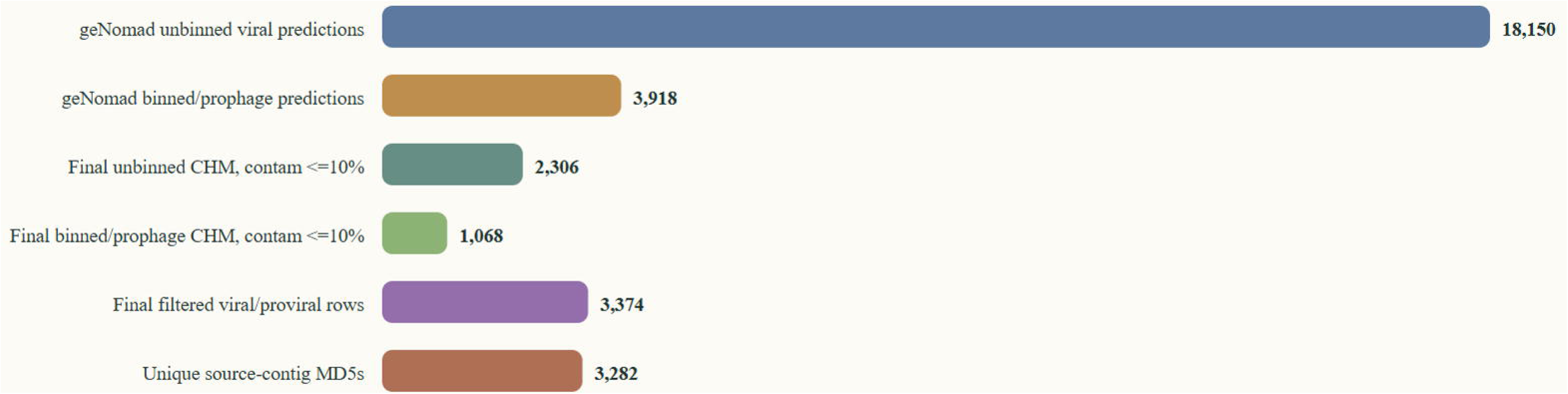
Viral and prophage candidate overview. Summary of unbinned/free-virus candidates and MAG-associated viral/prophage candidates generated from the final dog-first assembly and BASALT context. geNomad predicted 22,068 viral sequences, of which 3,374 candidate rows passed the final CheckV Complete, High-quality or Medium-quality (CHM) and contamination <= 10% criteria, representing 3,282 unique source-contig MD5s. Viral class-like assignments were dominated by *Caudoviricetes*. These records are candidate viral or proviral sequences and have not been dereplicated into vOTUs.

### Usage Notes

Users should select the DogMAG file layer that matches their analysis question. The 41 filtered dog-wise assembly FASTA files are the appropriate starting point for assembly-level remapping, contig-level annotation and independent viral discovery. Summary-level BASALT provenance and attrition metadata support workflow audit without requiring release of the complete transient BASALT workspace. The 3,418 reselected medium-quality-or-better MAG candidates and Supplementary Table 2 provide the pre-dereplication genome pool for users who want to apply alternative catalogue-unit definitions.

Together with the deposited GTDB-Tk-derived taxonomy table, the DogMAG 95% ANI representative collection can be configured as a minitax reference panel to generate rapid, detailed reference-based taxonomic composition profiles from canine gut metagenomes. This provides per-read recruitment and taxonomic-assignment summaries across the represented bacterial lineages. Resulting profiles should be interpreted relative to the taxonomic and genomic coverage of DogMAG rather than as exhaustive measurements of all organisms present in a sample.

Users should not treat BASALT bin selection as equivalent to a defined external ANI threshold. If species-level or strain-level units are required, users should rely on the external dereplication tables or rerun dereplication at their chosen thresholds using the deposited genome FASTA files and metadata. The 792 99% ANI representatives should be interpreted as the DogMAG strain-like comparison layer. The 135 95% ANI output should be interpreted as a species/SGB-like clustering layer, not as official SGB assignments.

When comparing DogMAG with other genome-resolved resources, users should match catalogue units carefully. The 99% ANI DogMAG layer can support strain-level catalogue context, whereas the 95% ANI layer provides species/SGB-like context; neither should be treated as directly equivalent to catalogues generated with different sample scopes, clustering thresholds, placement workflows or reporting conventions. External resources such as Branck et al. and Waltham should therefore be cited as complementary biological and methodological context rather than as strict numerical benchmarks [7,8].

The reference-panel validation tables should be used as a technical read-recruitment benchmark rather than as a substitute for biological community analysis. DogMAG recruitment in the mixed-kennel libraries is a source-cohort measure and may be favoured because those data contributed to catalogue construction. The Waltham subset is the independent validation layer. Database-specific taxonomic abundances, ecological differences between cohorts and biological associations are outside the scope of this Data Descriptor.

The unbinned viral candidate FASTA and table provide the cleaner free-virus discovery layer after final viral screening. MAG-associated viral/prophage candidates provide host-context and prophage-oriented records because these contigs are linked to BASALT bin/MAG candidates. Viral taxonomic assignments are reference-database-dependent annotations, especially at order, family and lower ranks. The viral results reported here are not dereplicated vOTUs; users requiring vOTU catalogues can perform viral dereplication on the deposited candidate sequences.

Previously published amplicon, short-read shotgun and source-cohort datasets linked to these samples are cited separately when reused for biological analyses. The DogMAG Data Descriptor is the citation target for the dog-wise assemblies, BASALT dog-first workflow outputs, final MAG representatives, viral/prophage candidates, metadata and workflow provenance described here.

## Supplementary Table Legends

**Supplementary Table 1. Dog-first bin/MAG attrition (20260727).** BASALT selected bin/version records, candidate versions, reselected medium-quality-or-better MAG candidates, high-completeness/low-contamination candidates, medium-only candidates and final externally dereplicated representatives, with counts removed at each filtering/dereplication step.

**Supplementary Table 2. MAG candidate and representative metadata (20260727).** Stable candidate and original-bin identifiers; selected and source FASTA names and portable paths; source origin and file checksums; binner or reassembly class; CheckM2 completeness and contamination; genome size, N50 and quality score; operational quality class; candidate counts per original bin group; dRep internal-filter, cluster and representative fields at 99% and 95% ANI; and GTDB-Tk classification fields where available. Blank dRep and GTDB-Tk fields identify the 196 candidates not retained by dRep’s internal filtering.

**Supplementary Table 3. Wet-lab provenance and DNA concentration metadata (20260727).** Source dataset, public sample identifier, sequencing platform, library name, extraction kit, library preparation workflow, DNA concentration, concentration unit and concentration source file for each sequencing library included in DogMAG.

**Supplementary Table 4. DogMAG assembly-to-read provenance manifest (20260727).** Public-facing table linking final dog-wise BASALT input assemblies to public read/sample identifiers, read role, source manifest, sequencing platform, ENA study accession and checksum fields where available.

**Supplementary Table 5. Viral/prophage candidate metadata (20260727).** Stable viral candidate identifier, source branch, source contig and assembly context, sequence MD5 where available, BASALT bin/MAG context, geNomad length, topology, score and taxonomy fields, and CheckV quality, completeness and contamination fields. Sequence files are organised by the stable candidate and source-contig identifiers in the associated article data package.

**Supplementary Table 6. Viral/prophage candidate quality summary (20260727).** Branch-level geNomad viral prediction counts, CheckV Complete/High/Medium counts after contamination filtering, unique source-contig MD5 counts and dominant broad viral taxonomy. These are not vOTU counts because viral dereplication was not performed.

**Supplementary Table 7. Balanced bacterial RefSeq-versus-DogMAG read-recruitment validation (20260727).** Paired per-library summary for 23 mixed-kennel source-cohort and 23 independent Waltham canine gut ONT metagenomes analysed against bacterial RefSeq and the DogMAG 95% ANI representative panel. Fields comprise validation library identifier, cohort, primary-read count, mapped-read fractions for each panel and their percentage-point difference, and final BestAln assigned-read fractions for each panel and their percentage-point difference. Detailed read-accounting and run-provenance files are included separately in the associated article data package.

Supplementary Table 8. Final dog-first assembly metrics and ENA deposition records (20260822). Assembly-level sequence metrics and ENA accessions for the 41 retained BASALT input assemblies, comprising 30 Flye long-read-only and 11 OPERA-MS hybrid assemblies. Fields include stable public assembly identifier, coassembly BioSample accession, primary-metagenome ERZ accession, ENA submission status, assembler and assembly class, linked FASTQ-record count, total assembly length, contig count, longest and shortest retained contigs, mean and median contig lengths, N50, N90, GC and N percentages, contig-length counts, and length-weighted BASALT total-average-depth estimate with source depth-table provenance. All sequence metrics were calculated after the 1,500-bp contig-length filter.

Supplementary Table 9. ENA deposition status of the 135 DogMAG ANI95 representatives (20260815). MAG alias, MAG-derived sample and BioSample accessions, NCBI and GTDB taxonomic metadata, source assembly and sample links, FASTA-derived contig count, genome-quality metrics, ERZ assembly-analysis accession where assigned, and assembly-submission status. The table distinguishes the 122 multi-contig MAG assemblies accepted by ENA and assigned ERZ accessions from the 13 single-contig MAG representatives retained in the DogMAG data package.

## Data Availability

Sample-linked input reads used for DogMAG assembly generation, binning, MAG recovery and validation are available from the European Nucleotide Archive under PRJEB75753 for Dog_M0 input data, PRJEB82125 for DMD input and provenance records, PRJEB85420 for CaniMeta/Serteperti input and provenance records, and PRJEB115259 for DogMAG sample registrations, newly deposited mixed-kennel long-read WGS data and associated DogMAG genome records. The PRJEB115259 FASTQ submission table uses public-coded sample, library and file identifiers rather than dog names. Previously published ENA datasets are cited as input-data and sample-provenance records. All 41 dog-wise primary metagenome assemblies are available as ENA analysis records ERZ29880033-ERZ29880073 and are cross-referenced in Supplementary Table 8. Of the 135 DogMAG ANI95 representative MAGs, all 122 multi-contig MAG assemblies were accepted by ENA as genome assembly analysis records and assigned ERZ accessions. Supplementary Table 9 lists these ERZ accessions together with the deposition status of all 135 representatives, including the 13 single-contig MAGs retained in the DogMAG data package. Reselected candidate MAG FASTAs, viral/prophage candidate sequences, metadata tables and detailed workflow outputs are organised in the associated article data package.

## Code Availability

Project-specific DogMAG scripts used for dog-first assembly generation, assembly collection, assembly preparation, BASALT input construction, read-to-assembly linkage, assembly metrics, final bin/MAG summarisation, viral candidate recovery, metadata generation, candidate re-selection, external dereplication, GTDB-Tk summarisation, reference-panel read-accounting and viral-result packaging are available from the DogMAG repository (https://github.com/Balays/DogMAG). The DogMAG repository also contains dataset-specific command lines, software versions and workflow-provenance notes for assembly generation, BASALT input construction, catalogue finalisation and data-package assembly.

The BASALT code version used for DogMAG is available at https://github.com/Balays/BASALT. BASALT was used as the core bin-comparison and BestBinset selection engine. The cleaned DogMAG publication snapshot is available at https://github.com/Balays/BASALT/tree/DogMAG-paper-v1.0. The generalized BASALT source changes are fixed at commit bdd4106a398db69bee83ea9ad1d331b00c041122 and documented in LARGE_RUN_PATCHES.md. These changes cover large-run correctness, performance, storage and resume behaviour, including restart-safe CheckM2 report validation, bounded reassembly resources, conservative cleanup, disk-space safeguards and resumable long-read intermediate handling. Dataset-specific rescue scripts, checkpoint edits, sample identifiers and private paths are intentionally excluded from the BASALT fork and remain documented with the DogMAG project workflow provenance.

Key DogMAG workflow scripts include:

scripts/assembly/run_flye_lr_only_dogfirst.sh

scripts/assembly/run_opera_ms_named_lrs_dogs.sh

scripts/assembly/collect_flye_lr_only_dogfirst_assemblies.sh

scripts/assembly/collect_hybrid_dogfirst_assemblies.sh

scripts/assembly/prepare_dogfirst_final_assemblies_for_basalt.sh

scripts/assembly/normalize_final_assemblies.py

scripts/assembly/link_reads_to_assemblies.py

scripts/assembly/assembly_metrics_fast.py

scripts/catalogue/summarize_basalt_binset.py

scripts/catalogue/run_catalog_unit_dereplication.py

scripts/catalogue/build_canmag_depletion_panels.sh

## Ethics statement

This study was conducted in accordance with applicable ethical and owner-consent requirements for canine faecal sample collection and data publication. Ethical approval was obtained from the Medical Research Council, Budapest, Hungary, under accession number BMEU/725-1/2022/EKU. Written informed consent was obtained from dog owners for sample collection and data publication.

## Author contributions

D.T. and Z.B. conceived and supervised the study. B.K. designed and implemented the computational workflow, dog-first assembly integration, BASALT analysis and manuscript data restructuring. N.J.Y., A.D. and T.J. contributed to sample processing, sequencing-library preparation, sequencing and source metadata curation. B.K. generated dog-wise assemblies, BASALT inputs, workflow provenance tables and downstream candidate-resource summaries. B.K. and D.T. drafted the manuscript. All authors reviewed and approved the final manuscript.

## Competing interests

The authors declare no competing interests.

## Funding

This study was supported by the Lendület I (Momentum I) Programme of the Hungarian Academy of Sciences (LP2020-8/2020) to D.T. and by the National Research, Development and Innovation Office (NKFIH OTKA FK 142676) to D.T. and (NKFIH OTKA K 142674 and NKFIH ADV 152705) to Z.B.

## Acknowledgements

We thank the dog owners, kennels and collaborators who provided canine faecal samples and source metadata for this resource. We thank the owners of the Dog_M0/Toti donor dog for their support in sample collection, Gabriella Kassai for providing samples from the Serteperti cohort, Ildikó Abonyi for providing samples from the Duna-menti Dumás cohort.

## Notes

### Competing Interest Statement

The authors have declared no competing interest.

